# Seesaw configuration of the ODC-OAZ1-AZIN1 axis governs sensitivity to polyamine-targeting inhibitors of breast cancer cells

**DOI:** 10.64898/2026.10.30.763251

**Authors:** Yuan Gao, Jinzhu Li, Yutong Jiao, Yan Ji, Mengxi He, Chuang Shi, Sen Liu

**Affiliations:** Cooperative Innovation Center of Industrial Fermentation (Ministry of Education & Hubei Province) & Key Laboratory of Fermentation Engineering (Ministry of Education), Wuhan 430068, China; Department of Pathology, The First Clinical Medical College of China Three Gorges University, Yichang Central People’s Hospital, Yichang 443000; Hubei Key Laboratory of Industrial Microbiology, National “111” Center for Cellular Regulation and Molecular Pharmaceutics, Hubei University of Technology, Wuhan 430068, China

**Author notes:** Corresponding Author: Sen Liu, Ph.D. These authors contributed equally to this work (Yuan Gao, Jinzhu Li, Yutong Jiao, Yan Ji).

**Keywords:** Protein-protein interaction, breast cancer, combination therapy, negative feedback loop, polyamine homeostasis

## Abstract

Polyamines are endogenous metabolites crucial for various cellular functions, and maintaining their cellular levels is critical for normal cell survival and fate. Polyamine homeostasis is managed by a complex metabolic network, wherein the ODC-OAZ1-AZIN1 protein-protein interaction axis plays a central role in regulating polyamine synthesis and membrane transport. While existing research has established that polyamine-targeted inhibitors (PTIs) significantly perturb this axis in various cancers, and that different breast cancer subtypes display distinct PTI sensitivities, the fundamental mechanism of drug sensitivity difference has remained poorly understood. Here, we combine large-scale bioinformatic analysis with comprehensive experimental verification to reveal a polarized seesaw configuration of the ODC-OAZ1-AZIN1 axis across diverse tumor types, including breast cancer. We show that this seesaw configuration is not only distinct across breast cancer subtypes but also associated with unique cellular states. Furthermore, we reveal that this configuration underpins the distinct sensitivity profiles of these subtypes to both single and combination treatments of PTIs. Crucially, we demonstrate that targeted re-configuration of this axis can shift the drug sensitivity of resistant cells. This study provides the first systematic mechanistic framework linking the configuration of the ODC-OAZ1-AZIN1 axis to PTI sensitivity, thereby providing a configuration-based framework for rational selection of PTI combination regimens and new personalized therapeutic strategies in breast cancer.

## 1. Introduction

Polyamines are ubiquitous polycationic metabolites with two or more amine groups. The representative polyamines in mammalian cells are spermidine, spermine, and their precursor putrescine. Positively charged under physiological conditions, most polyamines bind with negatively charged entities including DNA, RNA, and proteins, and are involved in diversified cellular functions, such as chromatin remodeling ^1^, gene transcription and translation^2^, and protein-protein interaction ^3^. Due to their indispensable biological roles, the cellular homeostasis of polyamines is strictly regulated by the polyamine metabolic network including polyamine synthesis, polyamine catabolism, and polyamine membrane transportation ^4^. In the polyamine synthesis pathway, L-ornithine is converted to putrescine by ODC (ornithine decarboxylase 1). ODC functions as a homodimer ^5^. Meanwhile, ODC can be inhibited by a protein inhibitor, the ODC antizyme 1 (OAZ1), whereas OAZ1 can be inhibited by a third protein, AZIN1 (OAZ1 inhibitor 1) ^6^. In addition, OAZ1 is negatively coupled with polyamine uptake ^7^, whereas the expression of AZIN1 positively correlated with the membrane import of extracellular polyamines ^8^. Ultimately, the ODC-OAZ1-AZIN1 axis functions as a feedback loop playing a central role in maintaining cellular polyamine homeostasis by orchestrating a tightly coordinated balance between polyamine synthesis and membrane transportation (Figure 1a).

**Figure 1.**
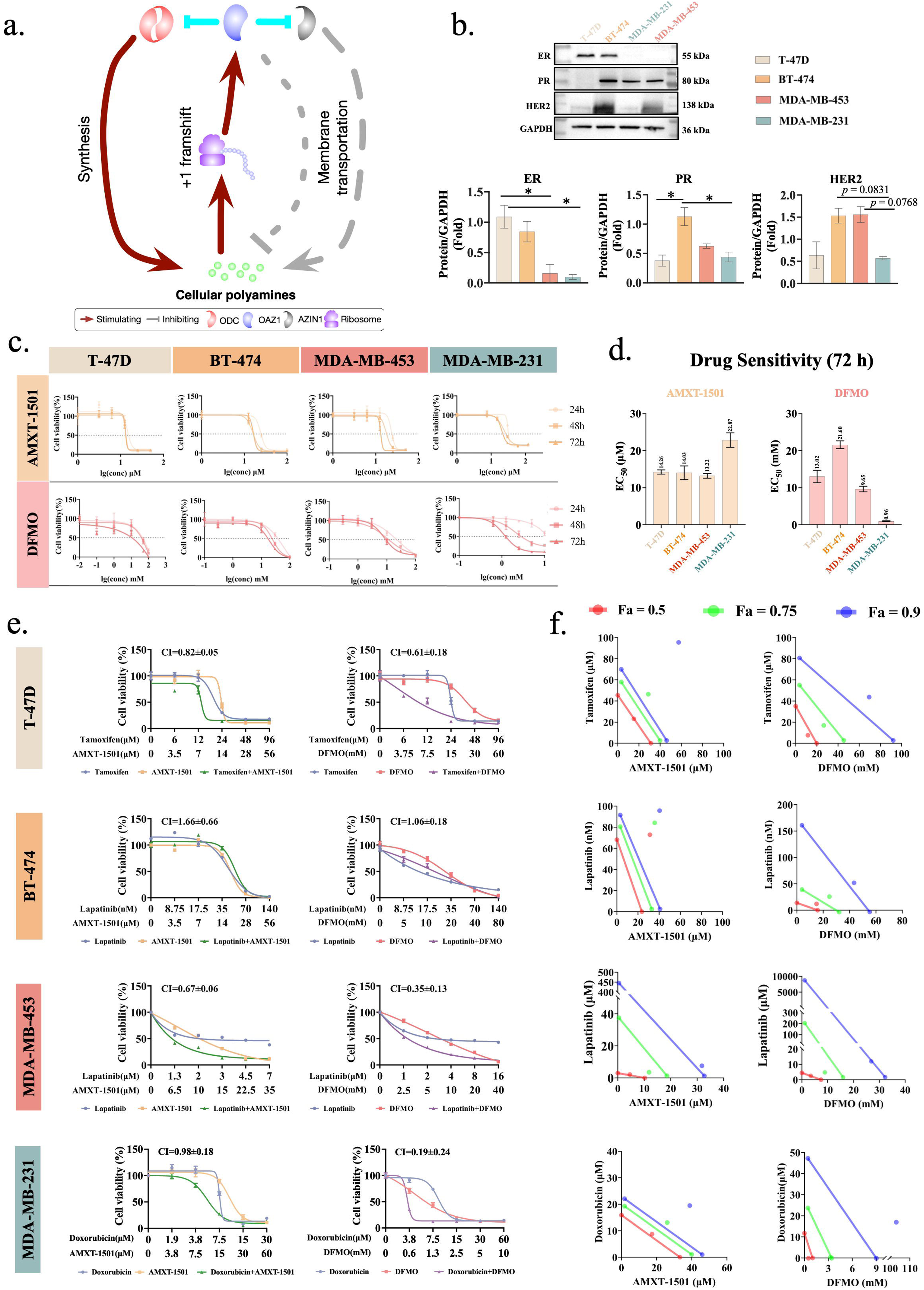
Breast cancer cells of different subtypes have varied sensitivity to PTIs. (a) A simplified scheme of the regulation of the ODC-OAZ1-AZIN1 axis. (b) Expression of the marker proteins (ER, PR, HER2) in the four breast cancer cell lines selected for investigation in this work. Quantitative data were from biological replications with error bars representing s.e.m. The *p* values were indicated as below throughout this manuscript: *, *p* ≤ 0.05; **, *p* ≤ 0.01; ***, *p* ≤ 0.001. (c, d) Sensitivity of four breast cancer cell lines to DFMO and AMXT-1501. EC_50_ values are calculated from the cell growth curves after the indicated treatment time (24 hours, 48 hours, 72 hours) and the data for 72 hours are shown in (d). (e, f) Cell growth curves of the four breast cancer cell lines under different combination treatments (e) and the corresponding normalized isobolograms (f). CI (combination index) values and the isobolograms were calculated with CompuSyn (ComboSyn, Inc.). CI values less than 1 indicates drug synergism, equal to 1 indicates additive effect, and larger than 1 indicates antagonism. F_a_ represents the fraction affected (the response) with 0 indicating 0% inhibition and 1 indicating 100% inhibition. A dot above (to the right of) the corresponding line indicates synergism and below indicates antagonism.

Dysregulation of polyamine homeostasis and the three proteins of the ODC-OAZ1-AZIN1 axis has been widely noticed in tumors ^9,10^. For example, polyamine levels were abnormally increased in tumor tissues and positively correlated with the histological grades of breast cancer ^11,12^. Similarly, elevated polyamine concentration was noticed in colorectal carcinoma and positively correlated with tumor stages ^13,14^. The expression level of ODC in human macrophages was upregulated in colitis-associated carcinogenesis, which impairs M1 macrophage activation ^15^. Increased levels of ODC and polyamines were detected in IL-33 induced M2-type macrophage differentiation and facilitated the migration and proliferation of esophageal squamous cell carcinoma (ESCC) cells ^16^. More generally, we noticed significantly increased ODC expression in more than ten kinds of tumors ^17^. On the contrary, being a negative regulator of polyamine uptake, OAZ1 is usually downregulated in cancer ^18^. In non-small cell lung cancer (NSCLC), for example, the patients with lower expression of OAZ1 had worse treatment response and poorer overall survival than the higher OAZ1 expression patients ^19^. Meanwhile, overexpressing OAZ1 reduced cell proliferation and inhibited forestomach tumor development ^20^. Conversely, the expression of AZIN1 is generally upregulated in different cancers including prostate cancer, breast cancer, lung cancer, liver cancer, and brain cancer ^21^.

Based on these facts, targeting the ODC-OAZ1-AZIN1 axis to perturb polyamine homeostasis (generally, reducing the polyamine level) in cancer cells becomes a promising strategy for cancer treatment ^17,22,23^. Since ODC is a rate-limiting enzyme for polyamine synthesis, inhibiting ODC’s activity is a sound option. There have been many clinical trials for evaluating the efficacy of DFMO (difluoromethylornithine), an irreversible inhibitor of ODC, in treating different cancers as a single drug (monotherapy), including neuroblastoma (NCT02395666, NCT04301843, NCT02679144), medulloblastoma (NCT04696029), prostate cancer (NCT00006101), cervical cancer (NCT00006079), gastric cancer (NCT02794428), bladder cancer (NCT00003814), non-melanomatous skin cancer (NCT00005884), and esophageal cancer (NCT00003076). Very recently, DFMO has been successfully approved to treat neuroblastoma. Although there are no inhibitors for OAZ1 and AZIN1, the uptake of extracellular polyamines can be blocked by polyamine transport inhibitors ^24^. AMXT-1501 is a well-studied polyamine transport inhibitor that can effectively inhibit polyamine uptake ^25^, and its single use showed varying effects on different cancer cells ^26–28^.

In targeted therapy, a disadvantage of monotherapy is that tumor cells could have counteracting compensatory regulations, so combination therapy with two or more drugs is generally more effective ^29^. For instance, when DFMO inhibits polyamine synthesis, tumor cells exhibit elevated uptake of extracellular polyamines in many cases ^30,31^, which hinders the clinical application of DFMO. For this reason, DFMO has been tested with other agents in multiple clinic trials for the treatment of various cancers (NCT02030964, NCT02139397, NCT01059071, NCT00983580, NCT01483144, NCT03794349, NCT01349881, NCT00033371, NCT00086736). Especially, the combination of DFMO and AMXT-1501 exhibited good synergistic effects ^25–28^, initiating clinical trials for a wide range of solid tumors including colorectal cancer, melanoma, ovary cancer, gastric cancer, esophageal cancer, breast cancer (NCT05500508, NCT03536728).

Besides compensatory regulations, cancer heterogeneity is another major reason of failed or compromised treatment outcomes in targeted therapy ^32,33^. The agents targeting the polyamine metabolic network also showed heterogeneity-dependent responses. For instance, on different human gastric cancer clones, the inhibiting effects of DFMO varied up to 460% ^34^. Extensive evidence has also emerged from breast cancer models. As a highly heterogeneous disease, breast cancer is usually classified into subtypes according to the expression levels of three marker proteins: ER (estrogen receptor), PR (progesterone receptor), and HER2 (human epidermal growth factor receptor 2). There are four typical breast cancer subtypes: luminal A (ER+, PR±, HER2-), luminal B (ER+, PR±, HER2+), HER2 positive (ER-, PR-, HER2+), and triple-negative (TNBC) or basal-like (ER-, PR-, HER2-) ^35^. Kremmer et al. ^36^ and Akinyele et al. ^37^ noticed that MCF-7 cells (ER+, PR+, HER2-; luminal A) were more sensitive to DFMO than MDA-MB-231 cells (ER-, PR-, HER2-; triple-negative). Similar heterogenous results were noticed in combination therapies even when polyamine-targeting inhibitors could sensitize breast cancer cells to anti-cancer drugs. Manni et al. ^38^ and Cohen et al. ^39^ proposed that the polyamine metabolic pathway could be involved in antiestrogen-mediated inhibition of breast cancer cells, and Cohen et al. ^40^ further verified that DFMO and 4-hydroxy-tamoxifen had an additive effect in inhibiting the growth of MCF-7 cells. Geck et al. ^41^ showed that DFMO increased the sensitivity of doxorubicin on triple-negative breast cancer (TNBC) cells MDA-MB-468 and SUM-159PT but not on non-TNBC cells. Akinyele et al. ^42^ found that DFMO and rapamycin, a mTOR pathway inhibitor, had additive efficacy on MCF-7 and MDA-MB-231 cells. Meanwhile, we demonstrated that simultaneously targeting ODC and A1 adenosine receptor had increased efficacies on breast cancer cells ^43^, although the extents of the beneficial effect varied on different cell lines.

Although the ODC-OAZ1-AZIN1 axis is critical for sustaining polyamine homeostasis, these proteins have largely been studied separately in previous studies. Consequently, it remains unknown how the integrated regulatory architecture of this axis is configured in different cancer cells, and whether and how such configuration relates to cellular heterogeneity. Therefore, in this work, using breast cancer cells of four clinical subtypes, we set out to investigate whether this axis is differently configured across these subtypes and how this configuration difference affects their heterogenous sensitivity to polyamine-targeting inhibitors (PTIs) and representative breast cancer therapies.

## 2. Results

### 2.1. Breast cancer cells of different subtypes have varied sensitivity to PTIs

First, to analyze the association between tumor heterogeneity and PTI sensitivity, we chose four representative breast cancer cell lines according to clinical breast cancer subtypes: MDA-MB-231 (ER-/PR-/HER2-; basal-like), T-47D (ER+/PR+/HER2-; luminal A), BT-474 (ER+/PR+/HER2+; luminal B), and MDA-MB-453 (ER-/PR-/HER2+; HER2+). The consistency between their subtypes and the expression levels of the three subtype markers (ER, PR, HER2) were confirmed (Figure 1b). We then tested the sensitivity of these cell lines to two representative PTIs: DFMO and AMXT-1501 (Figure 1c, 1d). MDA-MB-231 cells were less sensitive to AMXT-1501 than the other cell lines, but it had the highest sensitivity to DFMO. BT-474 cells had the lowest sensitivity to DFMO and moderate sensitivity to AMXT-1501, whereas MDA-MB-453 and T-47D cells showed moderate sensitivity to both DFMO and AMXT-1501. PTIs have long been known to be partially effective in cells and in vivo due to compensation mechanisms, and combination therapies of two or more drugs could have improved efficacy. Therefore, we investigated the synergistic potential of combining PTIs (DFMO or AMXT-1501) with representative first-line clinical breast cancer therapies: lapatinib (HER2 inhibitor), tamoxifen (ER inhibitor), and doxorubicin (a widely used chemotherapeutic agent for TNBC). The EC_50_ values of these clinical drugs on these four cell lines were obtained from the cell growth assay and used as the reference concentrations for combination treatment. According to the suggestion by Chou et al. ^44,45^, five step concentrations based on the EC_50_ values of these therapies were tested (Figure 1e), from which the normalized isobolograms at different combination ratios were derived (Figure 1f). Note that the goal of this study is not to directly compare the sensitivity of different cells to same treatments, so the clinical breast cancer therapies were only used on the corresponding subtype cells for best performance and potential clinical reference. To determine whether two therapies were synergistic or antagonistic in combination, we adopted the definition in the drug combination theory of Chou et al.^44,45^, and observed the following combination outcomes: (1) T-47D: AMXT-1501/tamoxifen, synergistic (CI < 1); DFMO/tamoxifen, synergistic (CI < 1); (2) BT-474: AMXT-1501/lapatinib, antagonistic (CI > 1); DFMO/lapatinib, antagonistic (CI > 1); (3) MDA-MB-453: AMXT-1501/lapatinib, synergistic (CI < 1); DFMO/lapatinib, synergistic (CI < 1); (4) MDA-MB-231: AMXT-1501/doxorubicin, synergistic (CI < 1); DFMO/doxorubicin, synergistic (CI < 1). Note that a drug pair was considered synergistic if at least one tested combination showed synergy.

Taken together, these data confirm that different breast cancer subtypes have different sensitivity to PTIs (DFMO and AMXT-1501). Meanwhile, these cells also respond differently to the combination treatment of PTIs and clinical breast cancer drugs.

### 2.2. Combination treatments perturb polyamine homeostasis in breast cancer cells similarly

Next, we investigated how different combinations of these inhibitors perturbed cellular polyamine homeostasis and whether the differences in drug sensitivity were driven by different changes in polyamine homeostasis. First, we quantified the basal polyamine (PA) levels of the four breast cancer cell lines (Figure 2a). MDA-MB-231 cells had the highest levels of both total PAs and individual PA species. MDA-MB-453 cells had high levels of putrescine (Put) and spermidine (Spd) but the lowest level of spermine (Spm). T-47D cells had the lowest Put level, but its Spm level was high. Then we quantified the cellular PA changes after the cells were treated for 72 hours by different inhibitors (Figure 2b). In all cell lines, the ODC inhibitor DFMO decreased the level of Put, the direct product of ODC. Except in MDA-MB-453 cells, DFMO also decreased the cellular Spd level. Altogether, the total PA levels in all cell lines were decreased by DFMO mono-treatment. In contrast, the mono-treatment of the polyamine transport inhibitor AMXT-1501 had little effect on the total PA levels in all cell lines, although it decreased the Put levels in T-47D cells and MDA-MB-453 cells. The mono-treatments of tamoxifen, lapatinib, and doxorubicin caused varied PA changes in these cells. In T-47D cells, tamoxifen decreased the Put level but had no significant effect on Spd, Spm, or total PA levels. In BT-474 cells, lapatinib caused significant increase in Put and the total PA level but did not affect Spd and Spm. In MDA-MB-453 cells, lapatinib increased the total PA level, mostly by the increase of Spd and Spm. In MDA-MB-231 cells, doxorubicin decreased all PAs. The PA changes affected by these three clinical breast cancer drugs suggest a significant regulatory crosstalk between their targets and the polyamine metabolic network, providing a rationale for their combination treatment with PTIs. In all combination treatments, the total PA levels were decreased in all cell lines. All separate PA species were also decreased by combination treatments, except that the Spm level in MDA-MB-453 did not change much.

**Figure 2.**
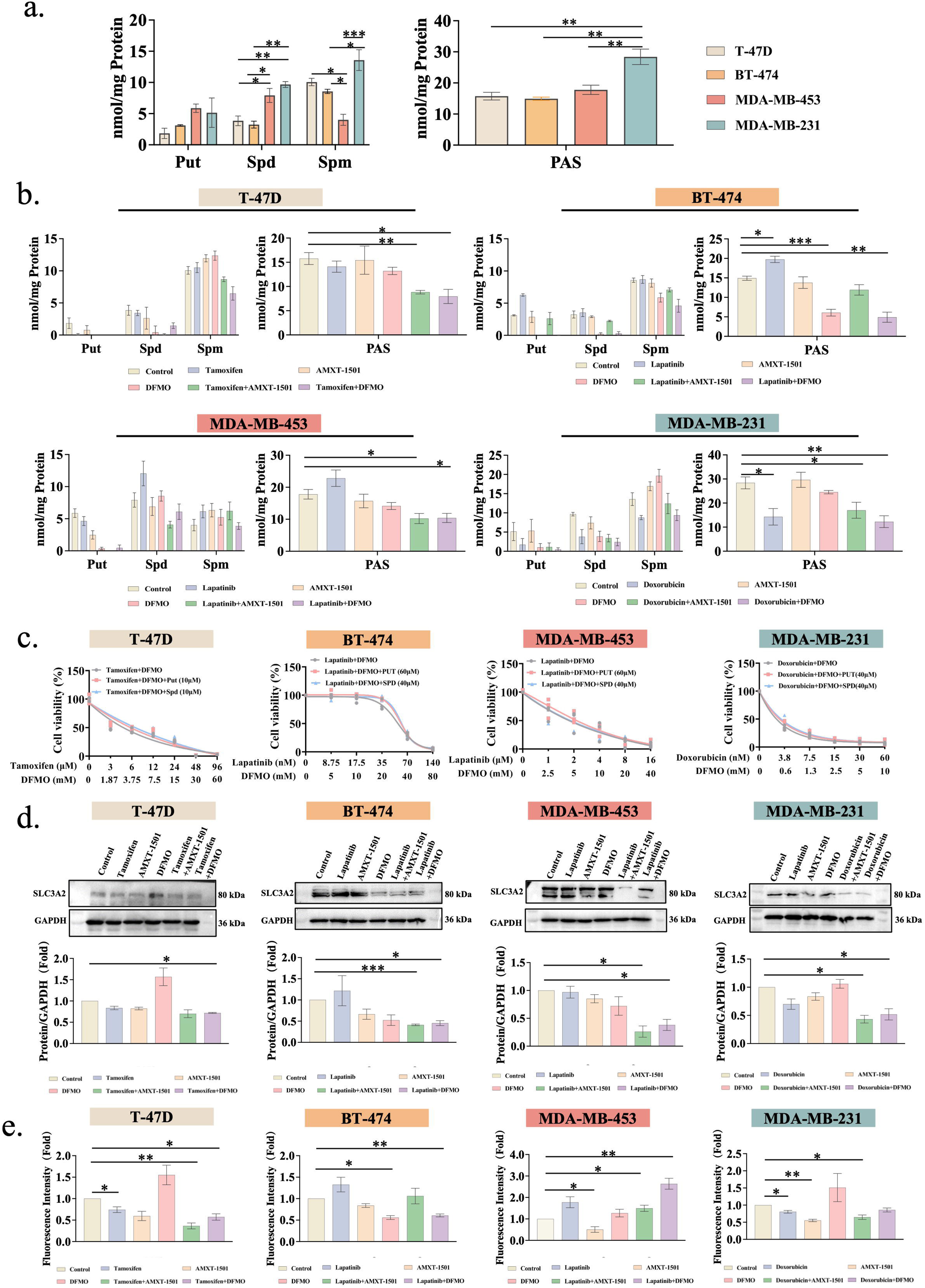
Combination treatments perturb polyamine homeostasis in breast cancer cells similarly. (a, b) Polyamine levels in four breast cancer cell lines under normal conditions (a) and different treatments for 72 hours (b). The concentrations of the inhibitors were determined from their EC_50_ values on different cell lines; concentrations used for T-47D cells were tamoxifen (15 μM), AMXT-1501 (10 μM), DFMO (10 mM), or their combinations; concentrations used for BT-474 cells were lapatinib (35 nM), AMXT-1501 (14 μM), DFMO (20 mM), or their combinations; concentrations used for MDA-MB-453 cells were lapatinib (3 μM), AMXT-1501 (15 μM), DFMO (10 mM), or their combinations; concentrations used for MDA-MB-231 cells were doxorubicin (15 nM), AMXT-1501 (30 μM), DFMO (2.6 mM), or their combinations. (c) Polyamine rescue assay results of four breast cancer cell lines under different treatments for 72 hours. (d) Changes of SLC3A2 levels in four breast cancer cell lines under different treatments for 72 hours. (e) Changes of the cellular fluorescent spermidine (MANT-Spd) of four breast cancer cell lines after different treatments for 72 hours. Data from biological replicates were shown in columns with error bars representing s.e.m.

Previous studies have demonstrated that if cell growth inhibition is caused by polyamine depletion, supplementing PAs in the culture media would rescue cell growth. The efficacy of this assay was confirmed on these four breast cancer cell lines via DFMO mono-treatment. When DFMO was combined with three clinical drugs, no cell growth recovery was noticed on all four cell lines by exogeneous PAs (Figure 2c). Because DFMO mono-treatment decreased the PA levels in all cell lines (Figure 2b) and it was known that DFMO treatment could induce PA uptake, we thought that under combination treatments, PA uptake in these cell lines might be blocked. Indeed, the polyamine transporter SLC3A2 was downregulated by all combination treatments in these four cell lines (Figure 2d). Mono-treatments had little effect on SLC3A2 in most cases with several exceptions: DFMO upregulated SLC3A2 in T-47D cells, both DFMO and AMXT-1501 downregulated SLC3A2 in BT-474 cells, and lapatinib upregulated SLC3A2 in BT-474 cells. Because PAs are also transported by transporters other than SLC3A2, we performed a polyamine uptake assay with a fluorescent spermidine (MANT-Spd) (Figure 2e). In all cell lines, AMXT-1501 mono-treatment inhibited the uptake of MANT-Spd. On the contrary, DFMO mono-treatment induced the increased uptake of MANT-Spd in T-47D, MDA-MB-453, and MDA-MB-231 cells, but the decreased uptake of MANT-Spd in BT-474 cells. However, the upregulation of SLC3A2 and the increased uptake of MANT-Spd by T-47D and MDA-MB-231 cells upon DFMO mono-treatment were consistent, as well as the downregulation of SLC3A2 and the decreased uptake of MANT-Spd in BT-474. Similarly, tamoxifen consistently induced downregulated SLC3A2 and MANT-Spd uptake in T-47D cells, as well as doxorubicin in MDA-MB-231 cells. Nonetheless, lapatinib majorly elevated SLC3A2 expression and MANT-Spd uptake in both BT-474 and MDA-MB-453 cells. In combination treatments, MANT-Spd uptake was inhibited in both T-47D and MDA-MB-231 cells, and lapatinib/DFMO inhibited MANT-Spd uptake in BT-474 cells. In other lapatinib combinations, the uptake of MANT-Spd was increased in both BT-474 and MDA-MB-453 cells.

Taken together, these data indicate that all PTI combinations perturbed the polyamine homeostasis similarly in these four breast cancer cell lines. Nonetheless, the polyamine homeostasis changes were not enough to explain the synergism/antagonism data of the combination treatments. Therefore, polyamine homeostasis perturbation appeared not be the differential factor of the PTI sensitivity across different breast cancer subtypes.

### 2.3. Configuration of the ODC-OAZ1-AZIN1 axis is polarized in breast cancer cells

The data above demonstrate that the combination treatments perturb polyamine levels and uptake similarly across all four cell lines, and exogenous polyamine supplementation fails to rescue growth under combination conditions, so the differential sensitivity cannot be attributed to the magnitude of polyamine depletion alone. We therefore hypothesized that the regulation network of polyamines might determine how each cell line responds to the same polyamine perturbation. Specifically, we turned our attention to the ODC-OAZ1-AZIN1 protein-protein interaction axis forming the feedback inhibition loop with polyamines (Figure 1a).

First, we wanted to know whether this axis is differently configured in clinical tumors. We collected and analyzed the gene copy-number alteration data of 37,744 tumor samples from 184 clinical tests including 36,487 patients from the cBioPortal database (http://cbioportal.org), among which 19,261 samples were profiled for *ODC*, *OAZ1*, and *AZIN1* copy-number alteration, and another 1,020 samples were profiled for *ODC* and *AZIN1* but not *OAZ1*. The definition of alteration types by this database was used, with deep deletion indicating homozygous deletion, shallow deletion indicating heterozygous deletion, low-level gain indicating few additional copies, and high-level gain (amplification) indicating more copies. Additionally, we reasoned that, as suggested by this database, deep-deletion and high-level gain are more biologically relevant. In both sample sets, *ODC* and *OAZ1* were steady (shallow deletion, diploid, or low-level gain), but *AZIN1* was significantly amplified (high-level gain) in significant portions. Gene copy-number alternation pattern analysis showed that *OAZ1* was steady in 18,994 of the 19,261 samples (98.6%), among which 17,553 samples (92.4%) were steady across these three genes. In the other 1,441 *OAZ1*-steady cases of the 18,994 samples with at least one of *ODC* and *AZIN1* showing significant alteration, the amplifications of *ODC* and *AZIN1* were largely misaligned: 1,261 cases (87.5%) were *AZIN1*-amplified and *ODC*-steady, whereas 128 cases (8.9%) were *ODC*-amplified and *AZIN1*-steady. This trend holds true in the 1,020-sample set as well. When classified into cancer types, ovarian epithelial tumor, breast cancer, hepatocellular carcinoma, bladder cancer, and prostate cancer were the five types with the top five cumulative amplification ratios in *ODC*, *OAZ1*, and *AZIN1* in these 37,744 samples (Figure 3a). Interestingly, four of these five cancer types are gender-associated. *AZIN1* had the most significant gene amplification overall and it had the highest gain ratio (12.4%) in breast cancer. Therefore, we then obtained gene copy-number data of 3,616 breast cancer samples from cBioPortal, among which 3,582 (99.1%) cases were *OAZ1*-steady (Figure 3b). The gene copy-number alteration patterns of these breast cancer samples are similar with that of the combined samples above (Figure 3c). Among these 3,582 cases, 2,937 cases (81.2%) were stable across all three genes, and 645 cases (18.8%) were *OAZ1*-stable but with significantly variations in *ODC* and/or *AZIN1*. In these 645 cases, the gain of *ODC* and *AZIN1* were misaligned in 637 cases (98.8%) as well (Figure 3d). Specifically, 94.0% (606) cases were *AZIN1*-amplified and *ODC*-stable, and 4.8% (31) were *ODC*-amplified and *AZIN1*-stable. Data from mRNA sequencing indicated that higher gene copy numbers generally result in higher gene expression levels for these three proteins.

**Figure 3.**
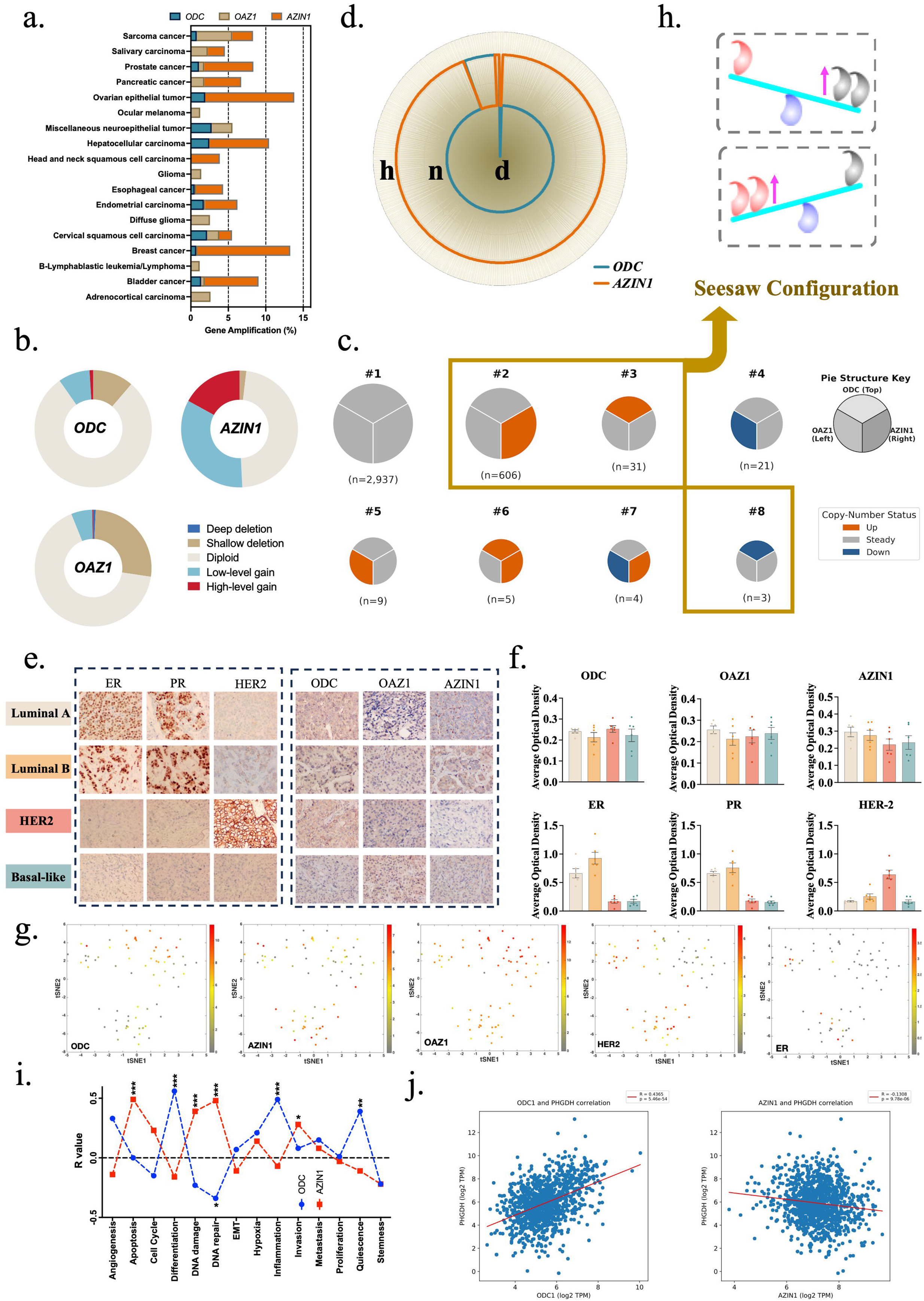
**Configuration of the ODC-OAZ1-AZIN1 axis is polarized in breast cancer cells**. (a) Classification of the 37,744 samples by cancer types and the percentages of the cases with significant copy number changes in *ODC*, *OAZ1*, and *AZIN1*. (b) Gene copy-number alteration data of 3,616 breast cancer samples from the cBioPortal database (http://cbioportal.org) profiled for *ODC*, *OAZ1*, and *AZIN1*. (c) Gene copy-number alteration patterns of the 3,616 breast cancer samples. Shallow deletion, diploid, and low-level gain were treated as steady. (d) Copy-number alteration of *ODC* and *AZIN1* in 645 *OAZ1*-steady breast cancer samples. Deep deletion and high-level gain were considered as significant. The outer circle (marked with **h)** indicates high-level gain, and the center (marked with **d**) indicates deep deletion. The middle circle (marked with **n**) between them represents stable level (shallow deletion, diploid, or low-level gain). The outer circle and the middle circle on the same radius belong to a same sample. For each gene, a broken in both the outer circle and the middle circle indicates deep deletion. (e, f) Immunohistochemistry (IHC) analysis of tumor tissues from clinical breast cancer patients. The ER, PR, and HER2 data matched clinical subtyping. Representative images are shown in (e), and statistical data are shown in (f). Data are shown as mean ± s.d., *n* = 6. (g) tSNE clustering of the scRNAseq data of the circulating tumor cells from clinical breast cancer patients. The expression levels of *HER2* (*ERBB2*), *ODC*, and *AZIN1* were shown according to the color scales. (h) Cartoon representation of the “seesaw configuration” of the ODC-OAZ1-AZIN1 axis in tumor cells. The pies in (c) within the solid frame indicates the seesaw configurations defined in this manuscript. (i) Correlation of the expression of *ODC* and *AZIN1* with different cellular functional states. For both (h) and (i), the original scRNAseq data were acquired by Jordan et al. ^46^ and the processed data were downloaded for analysis from CancerSEA ^50^. (j) Correlation between the expression levels of *PHGDH* and *ODC* or *AZIN1* in 1,135 OncoDB ^49^ clinical breast cancer samples.

Intrigued by the data analysis above, we collected clinical breast cancer samples for analysis. For each subtype, tumor samples from six patients were chosen. We analyzed the protein levels of the breast cancer subtype markers (HER2, ER, PR) and ODC/OAZ1/AZIN1 proteins (Figure 3e, Figure 3f). The relative expression of the marker proteins agreed well with their clinical subtyping. The expression of ODC, OAZ1, and AZIN1 had large variations without distinct patterns, possibly affected by the complexity and intratumor heterogeneity of clinical samples. Therefore, we resorted to single-cell mRNA sequencing (scRNAseq) data. Using two independent scRNAseq datasets from patient circulating tumor cells (CTCs) ^46,47^, we found that *OAZ1* had similar expression in all CTCs, but *AZIN1* and *ODC* showed mutually exclusive expression patterns (Figure 3g). Specifically, high *ODC* expression was observed in HER2-negative CTCs, whereas high *AZIN1* expression was in HER2-positive CTCs.

We define this configuration pattern of the ODC-OAZ1-AZIN1 axis as a ‘seesaw configuration’, characterized by the reciprocal, mutually exclusive upregulation patterns of ODC and AZIN1 against a steady OAZ1 baseline (Figure 3h). This configuration is also extendable to the mutually exclusive downregulation of ODC and AZIN1 leading to their similar relative changes. The correlation between this seesaw configuration and HER2 in breast cancer cells prompted us to think that, besides acting as compensating nodes for polyamine homeostasis, ODC and AZIN1 might be associated with different tumor states. Therefore, we analyzed the association of *ODC* and *AZIN1* expression with cellular signaling pathways using the scRNAseq data above. The associated cellular function states of *ODC* and *AZIN1* were mostly mutually exclusive (Figure 3i). ODC is positively associated with differentiation, inflammation, and quiescence, but negatively associated with DNA damage and DNA repair. On the contrary, AZIN1 was positively associated with apoptosis, DNA damage, DNA repair, and invasion, but negatively associated with differentiation and inflammation. To validate the mutually exclusive associations of ODC and AZIN1, the expression of *ODC* and *AZIN1* were evaluated against the expression of *PHGDH*, a gene not directly linked to polyamines but proved to negatively correlate with metastasis of breast cancer cells ^48^. The data of 1,135 clinical breast cancer samples from OncoDB ^49^ confirmed that *ODC* expression is positively correlated with *PHGDH*, whereas *AZIN1* expression negatively correlates with *PHGDH* (Figure 3j).

Collectively, these data demonstrate that in most tumor cases, the ODC-OAZ1-AZIN1 axis remains stable. But when it is perturbed, the ODC-OAZ1-AZIN1 axis exhibits a seesaw configuration in tumor cells including breast cancer cells. This seesaw configuration between ODC and AZIN1 is different from their dynamic compensation upon PTI treatments, because it is to fulfill basal requirements of different cellular function and states. The possible link of this basal configuration with breast cancer subtypes prompted us to hypothesize that this seesaw configuration might underlie the subtype-specific differences of PTI sensitivity observed above.

### 2.4. Seesaw configuration of the ODC-OAZ1-AZIN1 axis governs PTI sensitivity of breast cancer cells

We next investigated whether the seesaw configuration of the ODC-OAZ1-AZIN1 axis correlates with the PTI sensitivity of breast cancer cells of different subtypes. First, we quantified the cellular levels of these three proteins across the representative breast cancer cell lines, confirming that the axis is differently configured in each cell line (Figure 4a). While OAZ1 levels remained relatively stable across all cell lines, the expression of ODC and AZIN1 were negatively correlated (Figure 4b). Specifically, MDA-MB-231 and T-47D cells exhibited higher ODC expression, whereas BT-474 and MDA-MB-453 cells showed higher AZIN1 expression. Furthermore, we observed that ODC expression negatively correlated with PR (Figure 4c) and HER2 (Figure 4d), while AZIN1 expression positively correlated with PR (Figure 4e) and HER2 (Figure 4f). These expression patterns were also consistent with the observations above (Figure 3h).

**Figure 4.**
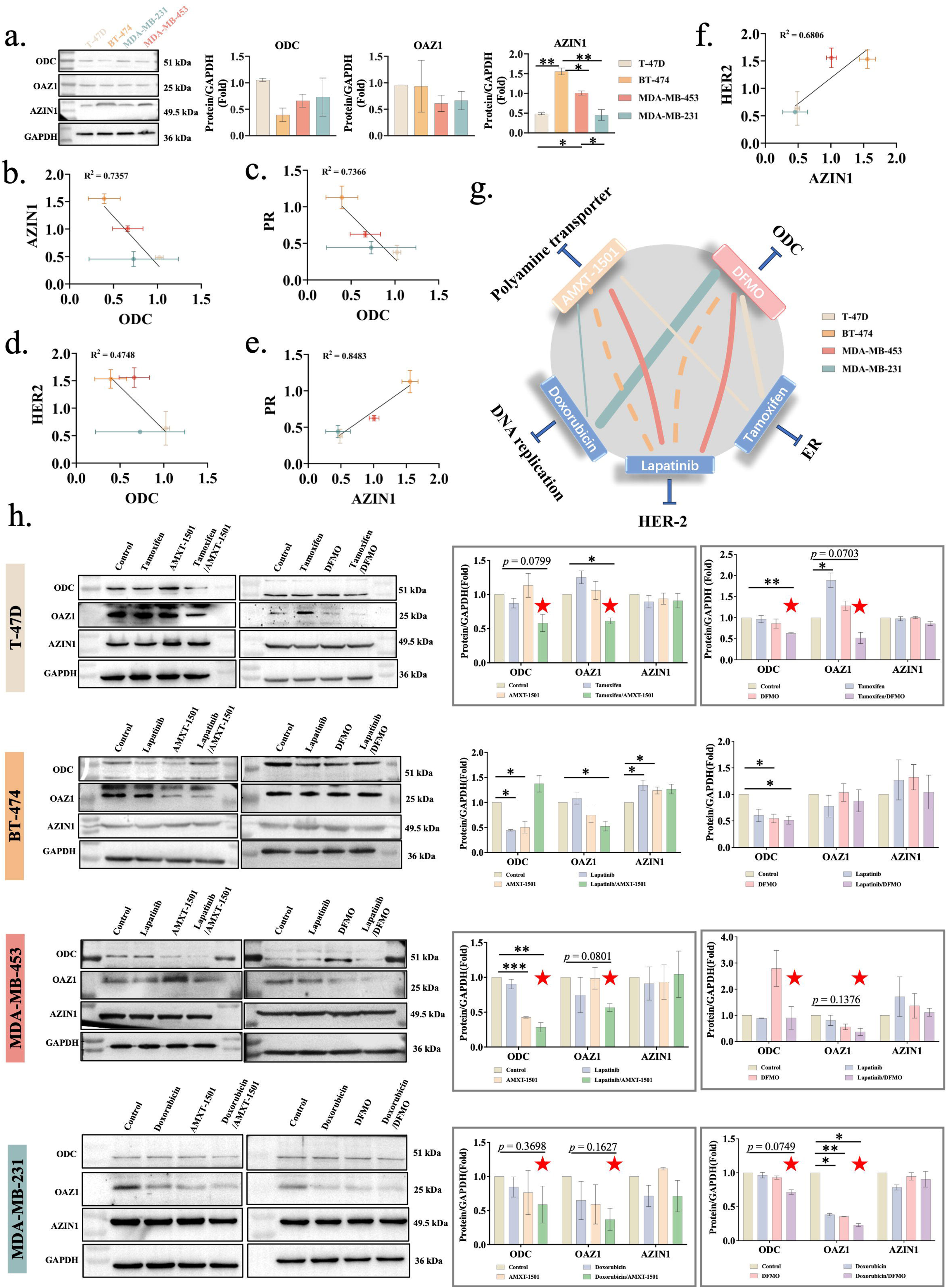
**Seesaw configuration of the ODC-OAZ1-AZIN1 axis governs PTI sensitivity of breast cancer cells**. (a) Expression of ODC, OAZ1, and AZIN1 in four breast cancer cell lines. Quantitative data from biological replicates are shown on the right. (b) Correlation between ODC expression and AZIN1 expression in four breast cancer cell lines. (c) Correlation between ODC expression and PR expression in four breast cancer cell lines. (d) Correlation between ODC expression and HER2 expression in four breast cancer cell lines. (e) Correlation between AZIN1 expression and PR expression in four breast cancer cell lines. (f) Correlation between AZIN1 expression and HER2 expression in four breast cancer cell lines. With four cell lines, (b)-(f) are only for demonstrating correlation trends. (g) Scheme of the effects of the combination treatments. Colors of the lines represent different cell lines. Solid lines indicate synergistic effect, and dashed lines indicate antagonistic effect. Width of the lines indicate the strength of the effect. (h) Expression of ODC, OAZ1, and AZIN1 under different monotreatment or combination treatment conditions in four breast cancer cell lines. The drug concentrations were identical to those used in Figure 2b, and the cells were treated for 72 hours, with quantitative data on the right. Solid borders are shown for combination treatments with synergistic effects. Data are shown as mean ± s.e.m.. Red stars indicate simultaneously down-regulated ODC and OAZ1.

We then assessed the relationship between these protein expression profiles and drug sensitivity. Previously, upon mono-treatment, intrinsic higher ODC levels paired with lower AZIN1 expression were associated with increased DFMO sensitivity; however, this pattern was associated with decreased sensitivity to AMXT-1501. T-47D cells represented an exception, exhibiting moderate sensitivity to both drugs despite their high ODC/low AZIN1 profile. In combination treatments, MDA-MB-453 cells (moderate ODC; moderate AZIN1) was sensitive to both DFMO/lapatinib and AMXT-1501/lapatinib. T-47D cells (high ODC; low AZIN1) showed high sensitivity to DFMO/tamoxifen but moderate sensitivity to AMXT-1501/tamoxifen. Similarly, MDA-MB-231 (moderate ODC; low AZIN1) was sensitive to DFMO/doxorubicin and slightly sensitive to AMXT-1501/Doxorubicin. Conversely, BT-474 (low ODC; high AZIN1) was antagonistic to DFMO/Lapatinib, but it was also insensitive to AMXT-1501/Lapatinib. Combining these data, the seesaw configuration of the ODC-OAZ1-AZIN1 axis represents a better explanation of the different drug sensitivity of these breast cancer cells (Figure 4g). To investigate the dynamic regulatory mechanism of combination treatments, we quantified changes in the axis following drug exposure (Figure 4h). Notably, in synergistic combinations, inhibitors induced a simultaneous downregulation of both ODC and OAZ1, whereas this coordinated downregulation was not observed in antagonistic combinations. Consistent with our seesaw model, AZIN1 remained stable when ODC decreased.

Collectively, these data confirm that the ODC-OAZ1-AZIN1 axis undergoes the polarized seesaw configuration in different breast cancer subtypes. Furthermore, the difference in the configuration of this axis also results in the sensitivity difference of these cancer cells to PTI treatments.

### 2.5. Reconfiguring the ODC-OAZ1-AZIN1 axis sensitizes T-47D cells to tamoxifen

The data above demonstrate that the basal configuration of the ODC-OAZ1-AZIN1 axis acts as a constraint on the direction of treatment-induced perturbation: cells configured toward the ODC pole are more vulnerable to ODC-directed perturbation, whereas cells configured toward the AZIN1 pole are buffered against it. Then, we continued to test whether deliberate reconfiguration of this basal axis shifts the sensitivity phenotype.

Noticing that the simultaneous down-regulation of ODC and OAZ1 coincided with the sensitivity to combination treatments in these four breast cancer cell lines, we investigated whether reconfiguring the ODC-OAZ1-AZIN1 axis by directly knocking down these two genes will shift their drug sensitivity. To recapture the effects of combination treatments, we performed siRNA-mediated knockdown of these two genes to the protein levels close to those observed in the combination treatments (Figure 4h). We note that siRNA-mediated knockdown and pharmacological inhibition are not mechanistically identical: DFMO inactivates ODC catalytic activity without reducing ODC protein abundance, whereas siRNA degrades ODC mRNA and reduces total protein. Nevertheless, both strategies are predicted to converge on reduced polyamine synthesis and, by extension, perturbation of the polyamine supply network. The seesaw model further predicts that whether this perturbation is sufficient to shift drug sensitivity depends on the basal configuration of the axis in each cell line.

When ODC and OAZ1 were knocked down together, significant decreases in Put, Spd, and total polyamines were observed in T-47D cells (Figure 5a). The polyamine transporter SLC3A2 only decreased in BT-474 and MDA-MB-453 cells (Figure 5b), but the membrane transport of MANT-Spd decreased across all cell lines (Figure 5c), indicating that polyamine membrane transport was downregulated by ODC/OAZ1 double knockdown. Accordingly, when the culture media were supplemented with exogenous PAs, the double knockdown also suppressed polyamine rescue except in T-47D cells (Figure 5d), indicating that the double knockdown did not fully inhibit polyamine membrane transport in this cell line. Finally, since ODC/OAZ1 double knockdown perturbs the polyamine metabolic network, its combination with breast cancer clinical drugs could show synergistical effects (increased sensitivity) as the combination drug treatments. Interestingly though, only T-47D cells were sensitized to tamoxifen by ODC/OAZ1 double knockdown (Figure 5e).

**Figure 5.**
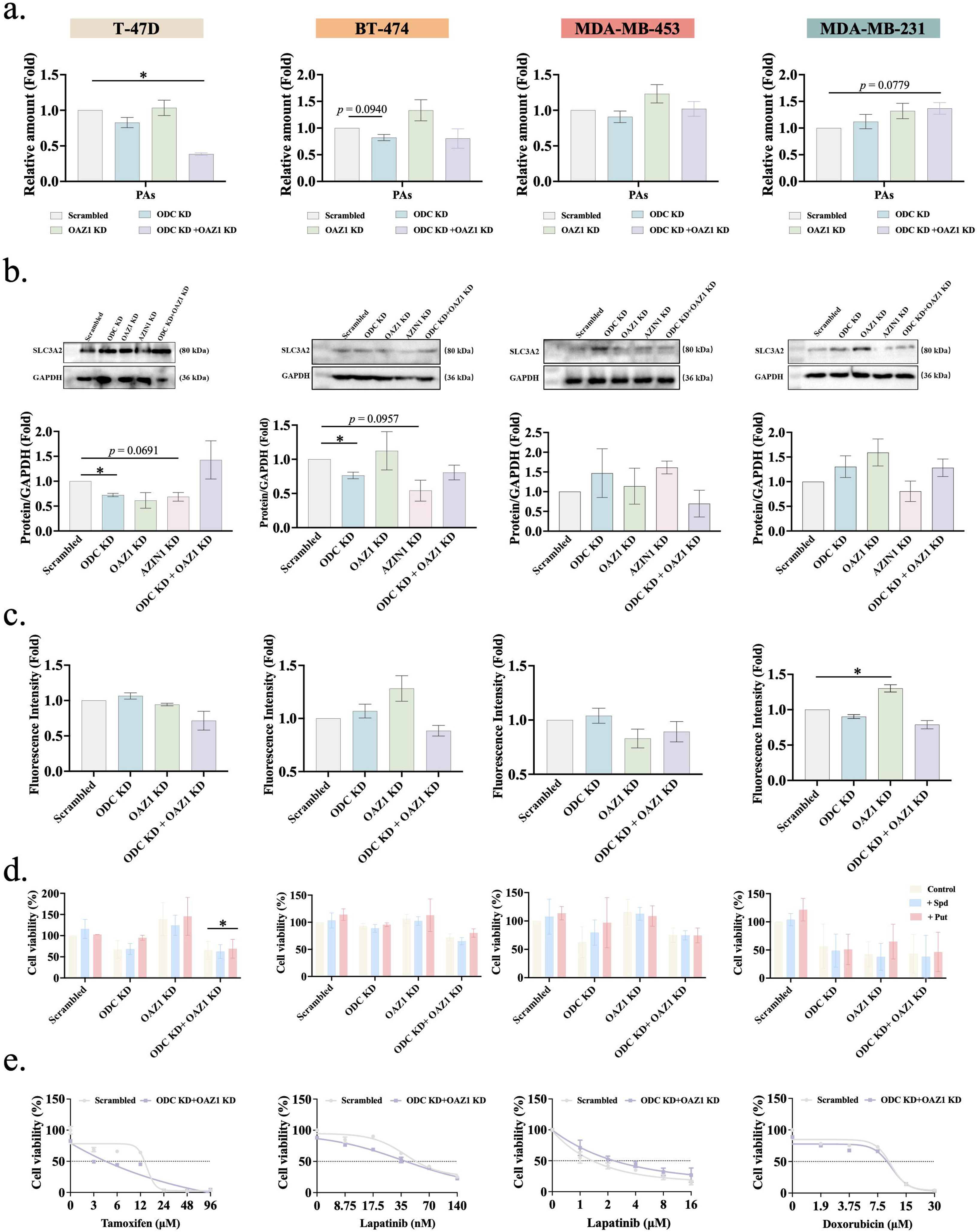
**Reconfiguring the ODC-OAZ1-AZIN1 axis sensitizes T-47D cells to tamoxifen**. (a) Changes of polyamine levels in four representative cell lines when *ODC* and *OAZ1* were knocked down together or alone. (b) Changes of SLC3A2 levels in four representative cell lines when *ODC* and *OAZ1* were knocked down together or alone. (c) Changes in the membrane transport of MANT-Spd in four representative cell lines when *ODC* and *OAZ1* were knocked down together or alone. (d) Changes of viability of four representative cell lines in the polyamine rescue assay when *ODC* and *OAZ1* were knocked down together or alone. The supplemented polyamine concentrations were same as those in Figure 2c. (e) Effects of *ODC*/*OAZ1* double knockdown on the sensitivity of four representative cell lines to indicated breast cancer clinical drugs.

Taken together, these data indicate that while the simultaneous decrease of ODC and OAZ1 protein levels mirrors the protein depletion observed in combination drug treatments, it was insufficient to sensitize all four cell lines. However, in T-47D cells, this double knockdown successfully recaptured the changes in polyamine homeostasis observed during drug combinations and sensitized the cells to tamoxifen. This selective sensitization of T-47D, but not the other three cell lines, is consistent with the seesaw model. T-47D cells are configured toward the high-ODC/low-AZIN1 pole, in which polyamine supply is dominated by ODC-driven de novo synthesis and the AZIN1-facilitated uptake arm is under-expressed. In this context, simultaneous reduction of ODC and OAZ1 disrupts the dominant supply route, and the weak uptake arm cannot compensate, as confirmed by the reduced SLC3A2 and MANT-Spd uptake and the failure of polyamine rescue. In drug combination treatment, DFMO enhances polyamine import, but tamoxifen downregulates SLC3A2 and suppresses MANT-Spd uptake, thereby blocking the compensation that DFMO induces. The net effect of the combination is a coordinated shutdown of both polyamine synthesis and the compensatory uptake response, producing the synergistic polyamine depletion and growth inhibition observed. In contrast, BT-474 cells (low ODC/high AZIN1) rely predominantly on AZIN1-facilitated uptake for polyamine supply; in these cells, ODC/OAZ1 knockdown disrupts the minor supply route and is buffered by the intact uptake arm, consistent with the lack of sensitization observed here and the requirement for AZIN1-targeted reconfiguration described in the next section. The absence of sensitization in MDA-MB-231 and MDA-MB-453 cells, in which both arms are moderately expressed, is likewise consistent with the model: the ODC/OAZ1 perturbation is partially compensated by the concurrent uptake capacity.

### 2.6. Reconfiguring the ODC-OAZ1-AZIN1 axis sensitizes BT-474 cells to PTI combinations

Given its characteristic resistance to both DFMO/lapatinib and AMXT-1501/lapatinib combination treatments, BT-474 cells (low ODC/high AZIN1) served as a critical model for testing whether reconfiguring the ODC-OAZ1-AZIN1 axis could affect PTI efficacy. The data above demonstrated that the double knockdown of *ODC* and *OAZ1* failed to sensitize this cell line, prompting us to investigate alternative regulatory mechanisms. Since the relative expression of ODC and AZIN1 were related to drug sensitivity, we compared the ODC/AZIN1 ratios of these four cell lines (Figure 6a) and noticed that BT-474 cells had the lowest ODC/AZIN1 ratio. Therefore, we hypothesized that elevating the ODC/AZIN1 ratio might sensitize BT-474 cells to PTIs.

**Figure 6.**
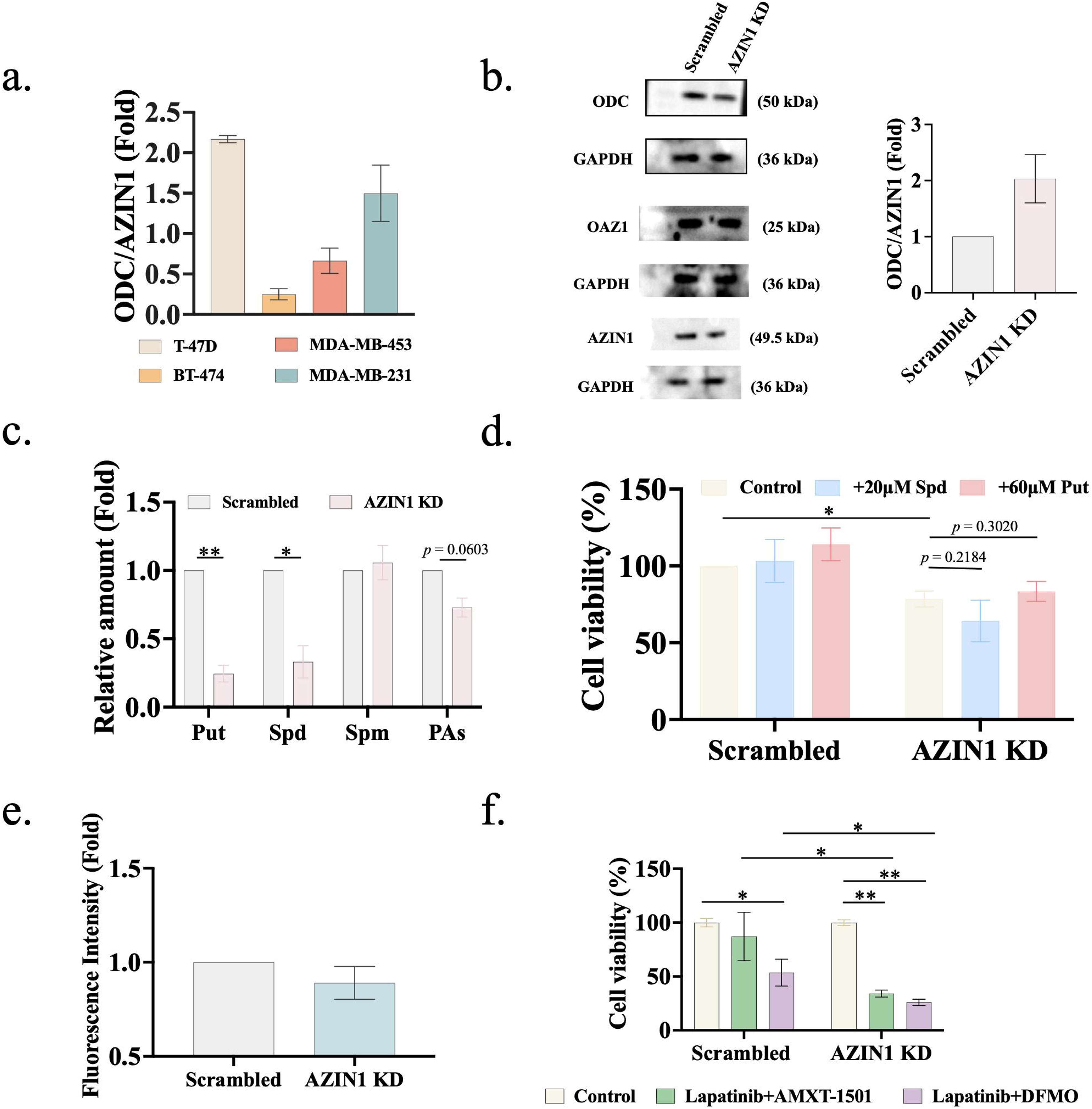
**Reconfiguring the ODC-OAZ1-AZIN1 axis sensitizes BT-474 cells to PTI combination treatments**. (a) Calculated ODC/AZIN1 ratio in four breast cancer cell lines. The expression data of ODC and AZIN1 were from Figure 4a. (b) Change of the expression of ODC, AZIN1, and their ratio in BT-474 when *AZIN1* was knocked down with siRNA. (c) Change of cellular polyamine levels upon *AZIN1* knockdown. (d) Change of cell viability from the polyamine rescue assay upon *AZIN1* knockdown. (e) Change of MANT-Spd uptake upon *AZIN1* knockdown. (f) Sensitivity of BT-474 cells to combination treatments after *AZIN1* knockdown. Data are shown as mean ± s.d. (*n* = 3).

Following siRNA-mediated knockdown of *AZIN1*, OAZ1 was stable and ODC was slightly decreased, possibly because that more OAZ1 could inhibit ODC and induce its degradation. However, AZIN1’s change was more significant, leading to the increase of the ODC/AZIN1 ratio (Figure 6b). This was accompanied by significant decrease in cellular putrescine and spermidine levels (Figure 6c). While spermine levels remained stable, the total polyamine level decreased (Figure 6c). Cell growth was inhibited upon *AZIN1* knockdown, but this effect could not be reversed by extracellular polyamine supplementation (Figure 6d). Consistently, *AZIN1* knockdown led to the down-regulation of the polyamine transporter SLC3A2 (Figure 5b) and the decrease of MANT-Spd uptake (Figure 6e). Finally, we observed that after increasing the ODC/AZIN1 ratio with *AZIN1* knockdown, the reconfigured BT-474 cells became more sensitive to both DFMO/lapatinib and AMXT-1501/lapatinib combination treatments (Figure 6f).

In conclusion, these data demonstrate that reconfiguring the ODC-OAZ1-AZIN1 axis in BT-474 cells by shifting the ODC/AZIN1 ratio changes its PTI sensitivity, confirming that the configuration of the ODC-OAZ1-AZIN1 axis governs PTI sensitivity in breast cancer cells.

## 3. Discussion and conclusions

Polyamines are essential endogenous metabolites required for a diverse array of cellular functions, including growth, proliferation, apoptosis, and aging. Maintaining polyamine homeostasis is critical for physiological stability. To this end, cells have evolved a complex metabolic network comprising key proteins such as ODC, OAZ1, and AZIN1. Dysregulation of the polyamine metabolic network is a hallmark of various pathologies, including inflammation, aging, and cancer. Notably, dysregulation of the polyamine metabolic network can directly cause polyaminopathies, a class of polyamine-related diseases including SRS (Snyder-Robinson syndrome), BABS (Bachmann-Bupp syndrome), and FABAS (Faundes-Banka syndrome). The clinical success of ODC inhibitors like DFMO in treating neuroblastoma and BABS syndrome underscores the potential of targeting the polyamine metabolic network for therapeutic intervention.

The ODC-OAZ1-AZIN1 axis plays critical role in regulating cellular polyamine homeostasis and its architecture and regulation has been well known; however, few previous reports have studied this axis as a whole quantitatively. In this work, we quantitatively investigated how the configuration of the ODC-OAZ1-AZIN1 axis affects tumor cells. Specifically, we revealed how the configuration of this axis in breast cancer cells correlates with differential sensitivity to PTIs.

In breast cancer, high levels of heterogeneity present a significant challenge to targeted therapies. While it is established that polyamine concentrations are elevated in many breast cancers compared to normal tissues, we and others have noticed distinct differences in polyamine regulation and drug sensitivity across different clinical subtypes of breast cancer ^36–43,51^. Nonetheless, the mechanism driving the observed variation in response to polyamine targeting inhibitors (PTIs) among different breast cancer subtypes has remained elusive.

In this study, we show that the drug sensitivity to PTIs of different breast cancer subtypes is not merely a function of absolute polyamine concentrations but is fundamentally governed by the intrinsic configuration of the ODC-OAZ1-AZIN1 axis. Combining large-scale bioinformatic analysis and extensive experimental validation, we reveal that breast cancer cells undergo a polarized “seesaw configuration” of this axis – a skewed and decoupled profile of the expression change of ODC and AZIN1 while OAZ1 remains stable. We show that this structural configuration in the polyamine metabolic network provides a mechanistic basis for the heterogeneity in PTI sensitivity observed across different breast cancer subtypes. Most significantly, we demonstrate that by reconfiguring this axis, we can shift the drug sensitivity of breast cancer cells.

Previous studies ^36,37,40,42^ have repeatedly observed that the luminal A breast cancer cell line MCF-7 was more sensitive to DFMO than the other breast cancer subtypes. Interestingly, the sole cell line showing consistent responses to drug combinations and ODC/OAZ1 double knockdown was a luminal A breast cancer cell line, T-47D. This convergence suggests that the polyamine metabolic network plays a more prominent role in this specific cell type, either through direct modulation of the subtype marker proteins (ER+/PR+/HER2-) or through broader, subtype-specific transcriptional programs.

Our data support that the seesaw configuration of the ODC-OAZ1-AZIN1 predicts the combination treatment potency of PTIs and clinical breast cancer targeted therapies, supporting the central role of this protein-protein interaction axis in regulating cellular polyamine homeostasis. The negative results from polyamine rescue and MANT-Spd uptake in the cell lines tested highlight that PTI combination treatments can efficiently block exogenous polyamine import through regulating the protein expression profile of this axis, thereby counteracting the compensatory polyamine-import upregulation induced by PTI mono-treatment. These data also suggest that the clinical drugs in these combinations are not pharmacologically inert bystanders but active modulators of the polyamine axis, whose effects on polyamine uptake are themselves configured by the seesaw state of the axis in each cell line.

It is important to note that the reconfiguration experiments in this work employed two distinct strategies: siRNA-mediated gene knockdown and pharmacological inhibition. These interventions are not mechanistically interchangeable. DFMO inhibits not only ODC but also other proteins, such as arginase ^52^, and AMXT-1501 is a broad inhibitor of polyamine membrane transporters and does not directly bind to AZIN1. Moreover, the proteins in the ODC-OAZ1-AZIN1 axis are also involved in pathways other than polyamine metabolism. For instance, OAZ1 binds cyclin D1 and induces its degradation ^53^, and hence AZIN1 may influence cyclin D1 stability ^54^. Therefore, the difference between the data from these two strategies across cell lines is not a contradiction but a reasonable outcome. Both approaches nonetheless converge on the same conclusion: the ODC-OAZ1-AZIN1 axis is a central regulator of polyamine homeostasis, and its protein-expression configuration is coupled to combination-drug sensitivity.

The seesaw configuration of the ODC-OAZ1-AZIN1 axis is not isolated to breast cancer. Our bioinformatic analysis identified ovarian epithelial tumor, breast cancer, hepatocellular carcinoma, bladder cancer, and prostate cancer as the five cancer types with the highest cumulative copy-number alteration ratios across ODC, OAZ1, and AZIN1. Four of these five are sex-hormone-associated malignancies, suggesting that the hormonal milieu may be a shaping force for the axis configuration. Indeed, estrogen and androgen signaling are known to cross-talk with polyamine metabolism in a cell-type-specific manner ^55–59^. However, broader association between this seesaw configuration and sex hormones requires future investigation.

Despite these insights, several limitations of this work warrant attention. First, the correlations between axis configuration and drug sensitivity are derived from four representative cell lines—one per major breast cancer subtype. With this small sample size, confidence intervals around the correlation coefficients are wide and statistical power to exclude moderate correlations is limited. We therefore treat the four-cell-line comparisons as a hypothesis-generating framework that identifies the seesaw configuration as the most intuitive explanation for the observed sensitivity pattern, rather than as a definitive statistical demonstration. Causal support for this model comes from the functional reconfiguration experiments in T-47D and BT-474, which demonstrate that deliberately shifting the axis configuration alters drug sensitivity. Second, protein and polyamine quantification relied on semi-quantitative methods (western blotting, HPLC, IHC), which limits the precision of cross-cell-line comparisons. Orthogonal quantification using MS-based proteomics and high-resolution metabolomics, ideally at single-cell resolution, would strengthen the quantitative basis of this framework. Third, the scRNAseq analysis draws on published circulating tumor cell (CTC) datasets from independent patient cohorts; while this independence is an advantage, the ODC/AZIN1 polarization observed may be influenced by disease stage, prior treatment, or CTC isolation methodology, and its robustness across more diverse cohorts remains to be confirmed. Fourth, most functional readouts were measured at a single endpoint (72 h), chosen because this time point captures robust PTI effects on polyamine levels in prior studies. The seesaw configuration is characterized here as a basal state; whether it is dynamic—shifting during the first 24–48 h of treatment, during chronic exposure, or as an acquired resistance mechanism—remains unresolved and represents a critical question for future work. Fifth, although we focus on the ODC-OAZ1-AZIN1 axis, hundreds of genes differ in expression across breast cancer subtypes, and the interplay between this axis and the broader transcriptional and signaling landscape of each subtype has not been fully addressed. Finally, validation in more physiologically relevant models—such as patient-derived xenografts (PDX), organoids, or in vivo combination-treatment studies—is essential to confirm the clinical translatability of these findings.

In conclusion, this work presents a first systematic explanation to the PTI sensitivity heterogeneity of breast cancer subtypes. By establishing the seesaw configuration of the ODC-OAZ1-AZIN1 axis as both a determinant and a manipulable target of PTI sensitivity, this work provides a framework for rational combination-therapy design and for re-sensitizing resistant tumor cells, with direct implications for precision oncology.

## Acknowledgements

We would like to thank the support from the other members of our lab. This research was funded by the National Natural Science Foundation of China [grant number 31670768, 31971150], the Department of Science and Technology of the Hubei Provincial People’s Government [grant numbers 2024AFA014, 2019CFA069].

## Author Contributions

Conceptualization, S.L.; methodology, Y.G., J.Z.L., Y.T.J., Y.J., M.X.H., and C.S.; validation, Y.G., J.Z.L., Y.T.J., Y.J., M.X.H., and C.S.; formal analysis, Y.G., J.Z.L., Y.T.J., Y.J., and S.L.; investigation, Y.G., J.Z.L., Y.T.J., Y.J., M.X.H., and C.S.; resources, S.L.; data curation, Y.G., J.Z.L., Y.T.J., and Y.J.; writing-original draft preparation, J.Z.L., Y.T.J., Y.J., and S.L.; writing-review and editing, S.L.; visualization, J.Z.L., Y.T.J., Y.J., and S.L.; supervision, S.L.; project administration, S.L.; funding acquisition, S.L. All authors have read and agreed to the published version of the manuscript.

## Data availability

Codes, as well as any materials used in this study, associated protocols and other supporting data, can be obtained from the corresponding author upon reasonable request.

## Materials and Methods

### Clinical ethics

This retrospective analysis of anonymized clinical data and biological samples was approved by the Medical Ethics Committee of Yichang Central People’s Hospital, with a waiver of informed consent.

### Database analysis

The cBioPortal (http://bioportal.org) data were acquired by querying the gene name *ODC1* (for ODC), *OAZ1*, *AZIN1*, and *ERBB2* (for HER2). For duplicated/overlapped cases, only the data from “TCGA, PanCancer Atlas” were kept. The data of cell lines were excluded. The samples had expression z-scores shown in “-2 (deep deletion)”, “-1 (shallow deletion)”, “0 (diploid)”, “1 (low-level gain)”, or “2 (high-level gain or amplification)”. The z-scores “-2” and “2” were considered as significant. In CancerSEA (http://biocc.hrbmu.edu.cn/CancerSEA), the single cell RNA sequencing data of Jordan et al. ^46^ and Aceto et al. ^47^ had the database IDs EXP0054 and EXP0055 respectively. The tumor cell classification was manually checked and verified according to the original reports. Pair-wise gene expression correlation analysis was performed on the OncoDB server (https://oncodb.org/).

### Immunohistochemistry (IHC) analysis

Clinical breast cancer samples were retrospectively selected based on clinical subtyping. Samples from six patients were selected for each subtypes. To confirm clinical subtypes, HER2, ER, and PR protein expression levels were evaluated along with ODC, OAZ1, and AZIN1. Formalin-fixed, paraffin-embedded tissue sections were baked at 75°C for 40 minutes, followed by deparaffinization using an environmentally friendly clearing agent (TO) at 70-80°C for 5 minutes. Antigen retrieval was performed via pressure-free boiling in an autoclave for 20 minutes, followed by rapid cooling under running water. To eliminate endogenous peroxidase activity, sections were incubated in a 3% hydrogen peroxide solution supplemented with 0.2% Tween-20 for 10-15 minutes and subsequently rinsed in phosphate-buffered saline (PBS) containing 0.2% Tween-20. Following section delineation with an IHC barrier pen, tissues were incubated at 37°C for 30 minutes with the following primary antibodies (with gentle agitation every 10 minutes to ensure uniform coverage): rabbit anti-AZIN1 polyclonal antibody (Proteintech, Cat. NO. 11548-1-AP), rabbit anti-OAZ1 polyclonal antibody (BBI, Cat. NO. D153218), rabbit anti-ODC polyclonal antibody (Affinity, Cat. NO. DF6712), rabbit anti-HER2 monoclonal antibody (VENTANA, Cat. NO. 790-2991), rabbit anti-ER monoclonal antibody (CONFIRM, Cat. NO. 790-4325), or rabbit anti-PR monoclonal antibody (CONFIRM, Cat. NO. 790-2223). Following two PBS washes, sections were incubated with an amplifier at 37°C for 15 minutes. After two additional PBS washes, sections were incubated with an HRP-conjugated goat anti-rabbit IgG secondary antibody (BBI, Cat. NO. D110058) at 37°C for 15 minutes. Following another round of PBS washes, signal visualization was achieved using 3,3’-diaminobenzidine (DAB) for 7 minutes, and the reaction was terminated with distilled water. Sections were counterstained, and blueing was performed using heated EDTA at 50°C. Pure water was used for all rinsing steps between EDTA retrieval and DAB development. Finally, the resulting IHC images were taken and then quantified using the IHC Image Analysis Toolbox in ImageJ.

### Cell Culture and Maintenance

The breast cancer cell lines T-47D, BT-474, and MDA-MB-231 were obtained from the China Center for Type Culture Collection (Wuhan, China); MDA-MB-453 cells were purchased from Wuhan Procell Life Science & Technology Co., Ltd (Wuhan, China). All cell lines were authenticated via cell viability monitoring, mycoplasma contamination testing, and short tandem repeat (STR) profiling. T-47D and BT-474 cells were cultured in RPMI 1640 medium (Hyclone, Cat. NO. SH30809.01) supplemented with 10% fetal bovine serum (FBS, Tocyto, Cat. NO. UT82901), 100 U/mL penicillin, and 100 µg/mL streptomycin (Hyclone, Cat. NO. SV30010) at 37 °C in a 5% CO_2_ incubator. MDA-MB-453 and MDA-MB-231 cells were cultured in Leibovitz L-15 medium (Hyclone, Cat. NO. SH30525.01) supplemented with 10% FBS, 100 U/mL penicillin, and 100 µg/mL streptomycin. All cells were used at <20 passages and routinely screened for mycoplasma.

### Cell Viability

Cell viability was quantified using the CCK-8 assay (Yeasen, Cat. NO. 40203ES88). Cells were seeded in 96-well plates and grown overnight. They were then treated with gradient dilutions of drugs for 24, 48, or 72 hours. After incubation, 5-10% CCK-8 reagent was added to each well for 2-4 hours at 37 °C, and absorbance was measured at 450 nm.

### Combination Treatment Analysis

The EC_50_ values of each tested drugs were obtained from mono-treatment. Then these values were used as the reference for combination treatment. Drug synergies were evaluated using the method established by Chou and Talalay ^44^. The Combination Index (CI), the isobologram, and Fa were calculated using CompuSyn software (ComboSyn, Inc., Paramus, NJ, USA), where CI < 1 indicates synergism, CI = 1 indicates an additive effect, and CI > 1 indicates antagonism.

### RNA Interference (siRNA) and Transfection

Cells were seeded in 6-well or 96-well plates and grown overnight to reach 70-90% confluence. Cells were transfected using Lipofectamine 2000 (Thermo Invitrogen, Cat. NO. 11668019) according to the manufacturer’s instructions. The following siRNA sequences were synthesized (Shanghai GenePharma Co., Ltd) and utilized: *ODC* (homo-793): 5’-GCGGAUUGCCACUGAUGAUTT-3’, 5’-AUCAUCAGUGGCAAUCCGCTT-3’; *OAZ1* (Homo-757): 5’-GCCUACACGUUCGAGAGAGTT-3’, 5’-CUCUCUCGAACGUGUAGGCTT-3’; *AZIN1* (homo-1073): 5’-CCAAGGUCUUACUACAUAUTT-3’, 5’-AUAUGUAGUAAGACCUUGGTT’; scrambled control: 5’-UUCUCCGAACGUGUCACGUTT-3’, 5’-ACGUGACACGUUCGGAGAATT-3’. Briefly, siRNA and transfection reagent were combined in serum-free media and incubated for 5 min at room temperature. The resulting DNA-lipid complexes were added to the cells and incubated for 6-8 hours before the medium was replaced with complete culture medium.

### Polyamine Quantification

Cells were washed with ice-cold PBS (Hyclone, Cat. NO. SH30256.01) and resuspended in 250 µL of lysis buffer (20 mM Tris-HCl, 150 mM NaCl, pH 8.0, 1% Triton X-100). After 10 min on ice, cells were centrifuged at 10,000 rpm for 10 min at 4 °C. The supernatant (200 µL) was collected, and total protein concentration was determined via BCA assay (Yeasen, Cat. NO. 20201ES76). To facilitate derivatization, the supernatant was treated with 20 µL of 50 µM 1,7-diaminoheptane (DAH, Sigma-Aldrich, Cat. NO. D17408), 500 µL of 2 mol/L NaOH, and 10 µL of benzoyl chloride. DAH served as an internal control. The mixture was vortexed for 30 s and incubated at 40 °C for 20 min. The reaction was terminated with 2.0 mL of saturated NaCl solution. Samples were extracted with dichloromethane, and the organic phase was dried overnight via volatilization. The precipitate was redissolved in 500 µL of methanol, filtered, and analyzed by LC-MS using an EclipsePlus C18 column (25 °C) under HESI positive ion mode.

### Polyamine Uptake Assay

Cells were seeded in 6-well plates and grown overnight, and then the medium was replaced with 2 mL/well of medium containing indicated drugs. After 72 hours of drug treatment, cells were incubated for additional 4 hours at 37°C with 1 mL/well of fresh medium containing 5 µM/L of MANT-Spd (synthesized by Wuxi AppTech Co., Ltd with > 95% purity). Then, cells were washed three times with 1 mM Spd-containing PBS (Hyclone, Cat. NO. SH30256.01). Cells were detached using trypsinization, resuspended in 1 mM Spd-containing PBS, and centrifuged at 1,500 rpm for 5 min. The cell pellet was lysed in RIPA buffer (Beyotime, Cat. NO. P0013B) for 10 min on ice and centrifuged at 10,000 rpm for 10 min. After BCA protein quantification, the supernatant was diluted in 500 µL ddH_2_O and transferred to a black-walled 96-well plate. Fluorescence was measured at an excitation of 341 nm and emission of 436 nm.

### Polyamine Rescue Assay

To determine if cell inhibition was caused by polyamine depletion, cells were treated with drugs for a specified duration. Specifically, cells were seeded in 96-well plates and grown overnight. Subsequently, the medium was replaced with complete medium supplemented with exogenous polyamines (Spd/Put) and indicated drugs. After 72 hours, cell viability was assessed via CCK-8 assay. Aminoguanidine (AG; MedChemExpress, Cat. NO. HY-B1041) was suggested to be supplemented to prevent the toxicity from the oxidation of exogenous polyamines ^60^, and we tested its proper concentration for each cell line. Specifically, the AG concentration was 0.5 mM for T-47D cells and 1.0 mM for MDA-MB-453 and MDA-MB-231. No AG was added for BT-474 cells due to obvious toxicity and neglectable toxicity from exogenouse polyamines.

### Western Blotting

Cells were washed with ice-cold PBS and lysed in radioimmunoprecipitation buffer (Beyotime, Cat. NO. P0013B) for 10 min on ice. Lysates were centrifuged at 10,000 rpm for 10 min at 4 °C, and protein concentration was measured via BCA assay. Equal amounts of protein were separated by SDS-PAGE and transferred to PVDF membranes (Millipore, Cat. NO. ISEQ00010). Membranes were blocked with 5% non-fat milk in TBST (Tris buffered saline with Tween-20) for 2 h at room temperature. Following overnight incubation with primary antibodies (anti-ODC rabbit polyclonal antibody: Affinity, Cat. NO. DF6712; anti-OAZ1 rabbit polyconal antibody: Thermo Invitrogen, Cat. NO. PA5-14385; anti-AZIN1 rabbit polyconal antibody: Proteintech, Cat. NO. 11548-1-AP; anti-SLC3A2 (CD98) rabbit monoclonal antibody: Thermo Invitrogen, Cat. NO. MA5-29574; anti-ER rabbit polyconal antibody: Affinity, Cat. NO. AF6058; anti-PR rabbit polyconal antibody: Affinity, Cat. NO. AF6106; anti-HER2 rabbit polyconal antibody: Affinity, Cat. NO. AF7681; anti-GAPDH rabbit polyconal antibody: Proteintech, Cat. NO. 10494-1-AP), membranes were incubated with HRP-conjugated secondary antibodies (Sangon Biotech, Cat. NO. D110058) for 2 h at room temperature. Proteins were visualized using ECL reagents (Boilogy, Cat. NO. BOLG001) and quantified via ImageJ densitometry, normalized to GAPDH.

## Statistical Analysis

Statistical analysis was performed using GraphPad Prism 9.5.0 or Gemini 3.6 Flash. Significant differences were evaluated using the Student’s t-test or the Tukey HSD post-hoc analysis for multiple comparisons. The *p* values were indicated as: *, *p* ≤ 0.05; **, *p* ≤ 0.01; ***, *p* ≤ 0.001.

## References

1. Imre, L. et al. Nucleosome destabilization by polyamines. Arch. Biochem. Biophys. 722, 109184 (2022).

2. Kaiser, A. & Agostinelli, E. Hypusinated EIF5A as a feasible drug target for Advanced Medicinal Therapies in the treatment of pathogenic parasites and therapy-resistant tumors. Amino Acids 54, 501–511 (2022).

3. Li, J. et al. Polyamines Disrupt the KaiABC Oscillator by Inducing Protein Denaturation. Molecules 24, 3351 (2019).

4. Gerner, E. W. & Meyskens, F. L. Polyamines and cancer: old molecules, new understanding. Nat. Rev. Cancer 4, 781–792 (2004).

5. Solano, F., Peñafiel, R., Solano, M. E. & Lozano, J. A. Equilibrium between active and inactive forms of rat liver ornithine decarboxylase mediated by L-ornithine and salts. FEBS Lett. 190, 324–328 (1985).

6. Kahana, C. Regulation of cellular polyamine levels and cellular proliferation by antizyme and antizyme inhibitor. Essays Biochem. 46, 47–62 (2009).

7. Mitchell, J. L., Judd, G. G., Bareyal-Leyser, A. & Ling, S. Y. Feedback repression of polyamine transport is mediated by antizyme in mammalian tissue-culture cells. Biochem. J. 299, 19–22 (1994).

8. Ramos-Molina, B., Lambertos, A. & Peñafiel, R. Antizyme Inhibitors in Polyamine Metabolism and Beyond: Physiopathological Implications. Med. Sci. 6, 89 (2018).

9. Thomas, T. & Thomas, T. J. Polyamine metabolism and cancer. J. Cell. Mol. Med. 7, 113–126 (2003).

10. Nakanishi, S. & Cleveland, J. L. Polyamine Homeostasis in Development and Disease. Med. Sci. 9, 28 (2021).

11. Kingsnorth, A. N., Wallace, H. M., Bundred, N. J. & Dixon, J. M. J. Polyamines in breast cancer. Br. J. Surg. 71, 352–356 (2005).

12. J, L., et al. Benefits of complete polyamine deprivation in hormone responsive and hormone resistant MCF-7 human breast adenocarcinoma in vivo. Anticancer Res. 20, (2000).

13. Weiss, T. S. et al. Polyamine levels of human colorectal adenocarcinomas are correlated with tumor stage and grade. Int. J. Colorectal Dis. 17, 381–387 (2002).

14. Linsalata, M. et al. Prognostic value of tissue polyamine levels in human colorectal carcinoma. Anticancer Res. 22, 2465–2469 (2002).

15. Singh, K. et al. Ornithine Decarboxylase in Macrophages Exacerbates Colitis and Promotes Colitis-Associated Colon Carcinogenesis by Impairing M1 Immune Responses. Cancer Res. 78, 4303–4315 (2018).

16. Mai, S. et al. Oesophageal squamous cell carcinoma–associated IL-33 rewires macrophage polarization towards M2 via activating ornithine decarboxylase. Cell Prolif. 54, e12960 (2020).

17. Chai, X. et al. The rational discovery of multipurpose inhibitors of the ornithine decarboxylase. FASEB J. 34, 10907–12921 (2020).

18. Coffino, P. Regulation of cellular polyamines by antizyme. Nat. Rev. Mol. Cell Biol. 2, 188–194 (2001).

19. Sun, Y. et al. Targeting HDAC/OAZ1 axis with a novel inhibitor effectively reverses cisplatin resistance in non-small cell lung cancer. Cell Death Dis. 10, 400 (2019).

20. Fong, L. Y. Y., Feith, D. J. & Pegg, A. E. Antizyme overexpression in transgenic mice reduces cell proliferation, increases apoptosis, and reduces N-nitrosomethylbenzylamine-induced forestomach carcinogenesis. Cancer Res. 63, 3945–3954 (2003).

21. Olsen, R. R. & Zetter, B. R. Evidence of a role for antizyme and antizyme inhibitor as regulators of human cancer. Mol. Cancer Res. 9, 1285–1293 (2011).

22. Meyskens, F. L., Jr. & Gerner, E. W. Development of difluoromethylornithine (DFMO) as a chemoprevention agent. Clin. Cancer Res. 5, 945–951 (1999).

23. Bassiri, H. et al. Translational development of difluoromethylornithine (DFMO) for the treatment of neuroblastoma. Transl. Pediatr. 4, (2015).

24. Muth, A. et al. Polyamine Transport Inhibitors: Design, Synthesis, and Combination Therapies with Difluoromethylornithine. J. Med. Chem. 57, 348–363 (2014).

25. Burns, M. R., Graminski, G. F., Weeks, R. S., Chen, Y. & O’Brien, T. G. Lipophilic lysine-spermine conjugates are potent polyamine transport inhibitors for use in combination with a polyamine biosynthesis inhibitor. J. Med. Chem. 52, 1983–1993 (2009).

26. Samal, K. et al. AMXT-1501, a novel polyamine transport inhibitor, synergizes with DFMO in inhibiting neuroblastoma cell proliferation by targeting both ornithine decarboxylase and polyamine transport: Synergistic Effect of DFMO/AMXT in NB. Int. J. Cancer 133, 1323– 1333 (2013).

27. Gamble, L. D. et al. Inhibition of polyamine synthesis and uptake reduces tumor progression and prolongs survival in mouse models of neuroblastoma. Sci. Transl. Med. 11, eaau1099 (2019).

28. Hayes, C. S. et al. Polyamine-Blocking Therapy Reverses Immunosuppression in the Tumor Microenvironment. Cancer Immunol. Res. 2, 274–285 (2014).

29. Jaaks, P. et al. Effective drug combinations in breast, colon and pancreatic cancer cells. Nature 603, 166–173 (2022).

30. Wu, R. et al. De novo synthesis and salvage pathway coordinately regulate polyamine homeostasis and determine T cell proliferation and function. Sci. Adv. 6, eabc4275 (2020).

31. Corral, M. & Wallace, H. M. Upregulation of Polyamine Transport in Human Colorectal Cancer Cells. Biomolecules 10, 499 (2020).

32. Marusyk, A., Janiszewska, M. & Polyak, K. Intratumor Heterogeneity: The Rosetta Stone of Therapy Resistance. Cancer Cell 37, 471–484 (2020).

33. Saunders, N. A. et al. Role of intratumoural heterogeneity in cancer drug resistance: molecular and clinical perspectives. EMBO Mol. Med. 4, 675–684 (2012).

34. Barranco, S. C., Ford, P. J. & Townsend, C. M. Heterogeneous survival responses of human gastric cancer clones to alpha difluoromethylornithine in vitro. Invest. New Drugs 4, 337–345 (1986).

35. Holliday, D. L. & Speirs, V. Choosing the right cell line for breast cancer research. Breast Cancer Res. 13, 215 (2011).

36. Kremmer, T. et al. Comparative studies on the polyamine metabolism and DFMO treatment of MCF-7 and MDA-MB-231 breast cancer cell lines and xenografts. Anticancer Res. 11, 1807– 1813 (1991).

37. Akinyele, O. & Wallace, H. M. Characterising the response of human breast cancer cells to polyamine modulation. Biomolecules 11, 743 (2021).

38. Manni, A., Wright, C., Demers, L. & Luk, G. Polyamines and hormonal regulation of N-nitrosomethylurea-induced rat mammary tumor growth in vivo. Cancer Res. 46, 4938–4941 (1986).

39. Cohen, F. J., Manni, A., Glikman, P., Bartholomew, M. & Demers, L. Involvement of the polyamine pathway in antiestrogen-induced growth inhibition of human breast cancer. Cancer Res. 48, 6819–6825 (1988).

40. Cohen, F. J., Manni, A., Glikman, P., Bartholomew, M. & Demers, L. Interactions between growth factor secretion and polyamines in MCF-7 Breast Cancer Cells. Eur. J. Cancer Clin. Oncol. 26, 603–608 (1990).

41. Geck, R. C. et al. Inhibition of the polyamine synthesis enzyme ornithine decarboxylase sensitizes triple-negative breast cancer cells to cytotoxic chemotherapy. J. Biol. Chem. 295, 6263–6277 (2020).

42. Akinyele, O. & Wallace, H. M. Understanding the Polyamine and mTOR Pathway Interaction in Breast Cancer Cell Growth. Med. Sci. 10, 51 (2022).

43. Ma, H. et al. Dual inhibition of ornithine decarboxylase and a1 adenosine receptor efficiently suppresses breast tumor cells. Front. Oncol. 11, 636373 (2021).

44. Chou, T.-C. Theoretical Basis, Experimental Design, and Computerized Simulation of Synergism and Antagonism in Drug Combination Studies. Pharmacol. Rev. 58, 621–681 (2006).

45. Chou, T.-C. Drug Combination Studies and Their Synergy Quantification Using the Chou-Talalay Method. Cancer Res. 70, 440–446 (2010).

46. Jordan, N. V. et al. HER2 expression identifies dynamic functional states within circulating breast cancer cells. Nature 537, 102–106 (2016).

47. Aceto, N. et al. AR Expression in Breast Cancer CTCs Associates with Bone Metastases. Mol. Cancer Res. 16, 720–727 (2018).

48. Rossi, M. et al. PHGDH heterogeneity potentiates cancer cell dissemination and metastasis. Nature 605, 747–753 (2022).

49. Tang, G., Cho, M. & Wang, X. OncoDB: an interactive online database for analysis of gene expression and viral infection in cancer. Nucleic Acids Res. 50, D1334–D1339 (2022).

50. Yuan, H. et al. CancerSEA: a cancer single-cell state atlas. Nucleic Acids Res. 47, D900–D908 (2019).

51. Song, Qi, Wang, Yixuan, & Liu, Sen. Subtype-specific transcription factors affects polyamine metabolism and tumor microenvironment in breast cancer. Cancer Innov. (2025) doi:10.1002/cai2.138.

52. Selamnia, M., Mayeur, C., Robert, V. & Blachier, F. α-Difluoromethylornithine (DFMO) as a potent arginase activity inhibitor in human colon carcinoma cells. Biochem. Pharmacol. 55, 1241–1245 (1998).

53. Newman, R. M. et al. Antizyme targets cyclin D1 for degradation. J. Biol. Chem. 279, 41504– 41511 (2004).

54. Kim, S. W. et al. Regulation of cell proliferation by the antizyme inhibitor: evidence for an antizyme-independent mechanism. J. Cell Sci. 119, 2583–2591 (2006).

55. Kumar, R. et al. Androgen receptor drives polyamine synthesis, creating a vulnerability for prostate cancer. Cancer Res. 86, 1148–1165 (2026).

56. Shukla-Dave, A. et al. Ornithine Decarboxylase Is Sufficient for Prostate Tumorigenesis via Androgen Receptor Signaling. Am. J. Pathol. 186, 3131–3145 (2016).

57. Poirier, D., Roy, J. & Maltais, R. A Targeted-Covalent Inhibitor of 17β-HSD1 Blocks Two Estrogen-Biosynthesis Pathways: In Vitro (Metabolism) and In Vivo (Xenograft) Studies in T-47D Breast Cancer Models. Cancers 13, 1841 (2021).

58. Zhu, D. et al. Single-Cell Transcriptome Analysis Reveals Estrogen Signaling Coordinately Augments One-Carbon, Polyamine, and Purine Synthesis in Breast Cancer. Cell Rep. 25, 2285–2298.e4 (2018).

59. Zhu, Q., Jin, L., Casero, R. A., Davidson, N. E. & Huang, Y. Role of ornithine decarboxylase in regulation of estrogen receptor alpha expression and growth in human breast cancer cells. Breast Cancer Res. Treat. 136, 57–66 (2012).

60. Holbert, C. E. et al. Autophagy induction by exogenous polyamines is an artifact of bovine serum amine oxidase activity in culture serum. J. Biol. Chem. 295, 9061–9068 (2020).

